# Diurnal butterfly diversity in a human-modified landscape of the subtropical montane forest of NW Argentina

**DOI:** 10.1101/2024.05.14.592973

**Authors:** Andrea del V. Guanuco, Mariano Ordano, Laura C. Pereyra, María José Barrionuevo, Noelia V. Gonzalez Baffa Trasci, Marcos Vaira

## Abstract

The change and intensification in land-use are currently among the main causes of species declines and local extinctions around the world. Therefore, forecasting changes in species diversity concerning habitat conditions may be crucial for conservation strategies. We explored diurnal lepidopteran diversity in a modified landscape of subtropical montane forests of Jujuy, NW Argentina. We considered that degradation of the natural forest habitat would likely impact resources crucial for butterflies, consequently altering both species richness and composition within these forests. We assessed and compared alpha diversity through Hill numbers diversity profiles and beta diversity through the beta-diversity partitioning. Additionally, we employed a permutational multivariate analysis of variance, and rank-abundance curves of butterfly species across different habitat types. Our results suggest that land-use changes diminish the number of forest-dependent species and increase species more tolerant to modified habitats and open areas. While alpha diversity did not decrease as land use changed, beta diversity showed significant changes in butterfly species composition, with a worrying reduction of forest-related species in altered habitats. Species composition became increasingly dominated by open area butterfly species resulting in biota homogenization, with potential consequences for ecosystem functioning and services in these forests. Further research on the mechanisms underlying the effects of human-induced habitat changes on forest butterfly diversity could help clarify which mitigation strategies are most likely to be successful for the conservation of butterflies of the subtropical montane forests.

## Introduction

The transformation of large areas of native forests into mosaics of human-modified habitats with varying land-uses is a major driver of biodiversity loss, posing an increasing threat to species extinctions worldwide (Newbold et al. 2015; Picanco et al. 2017). As habitat deterioration accelerates, predicting shifts in species diversity has become essential for effective conservation strategies. Several studies have highlighted the decline of various species, including butterflies, in response to habitat loss and fragmentation caused by human activities (Steffan-Dewenter & Tscharntke 2000; Uehara-Prado et al. 2007; Vasconcelos et al. 2019). Species communities respond to these changing conditions in different ways, with some managing to persist and others facing extinction as habitats shrink or become isolated (Barlow et al. 2009; Thomas 2016; Vasconcelos et al. 2019).

Among insect groups, butterflies have been identified as particularly effective models for detecting responses to land-use changes (Barlow et al. 2007a; Lazzeri et al. 2011; Sagwe et al. 2015; Orta et al. 2022). This is because they are highly sensitive and respond quickly to alterations in habitat conditions, such as temperature, microclimate, humidity, and lighting, as well as changes in vegetation composition and structure (Thomas et al. 2004; Thomas 2005; Bonebrake et al. 2010; Checa et al. 2014; Barlow et al. 2007b; Uehara-Prado & Freitas 2007). Moreover, the relatively well-documented systematics and taxonomy of Lepidoptera further enhance their utility as indicators in ecological studies (Thomas 2005; Lucci Freitas et al. 2014). Accordingly, several studies have shown that butterflies exhibit marked responses to land-use change, with distinct effects on species richness and composition. These include local extinctions or reduced persistence of certain threatened or rare species, and increases in the abundance of more non-threatened or widespread species (Barlow *et al*. 2007ab; Bubová *et al*., 2015; Thomas, 2016; Vasconcelos *et al*. 2019). These responses have been observed not only at the family level but also within closely related groups, including species from the same subfamily (Veddeler *et al*. 2005).

The Southern Andean Yungas ecoregion in Argentina, defined as an outstandingly biodiverse region, is considered a critical area for biodiversity conservation because of the high levels of species richness and endemism that are among the highest in the country, but also due to increasing land-use changes that threaten several species (Brown *et al*. 2006). This ecoregion covers only 2% of the country’s land and is the second most diverse ecoregion (Pacheco & Brown 2006; Nanni *et al*. 2020) that extends as a narrow strip of montane forest in the NW region of the country, corresponding to the southernmost extension of Neotropical cloud forests (Kapelle & Brown 2001). The Southern Andean Yungas ecoregion stands out for its high heterogeneity of environments with important extensions of subtropical montane forests, small plots with subsistence agriculture, logging sectors, large areas with commercial crops, and areas with different degrees of urbanization (Malizia *et al*. 2012). In such human-intervened landscapes, it is particularly relevant to investigate the response of local biodiversity to human pressures and to estimate biodiversity changes at local scales to determine conservation priorities (Reynolds *et al*. 2011). However, the lack of information on how species respond to landscape heterogeneity and land-use change in these montane forests poses a major challenge for reversing species declines and for the preservation and restoration of suitable habitats to maintain species diversity (Izquierdo & Grau 2009). Successful long-term conservation plans will require a better understanding of the relationships between land-use intensification and species composition to identify key drivers of species diversity in the region.

In this study, we describe diurnal lepidopteran diversity in a portion of the subtropical montane forests of Jujuy, NW Argentina where human intervention has transformed the entire area into a mosaic formed by patches of either actively used land (*Eucalyptus* plantations), degraded but currently unused land (secondary grasslands) and remnants of native forest. We expect that butterfly diversity in the area should be particularly sensitive to human-induced changes, showing the highest richness and densities within patches of native forests and reduced diversity within more altered patches.

## Material and methods

### Study area

The study was conducted at Finca “Las Capillas”, (24° 05LJ 34’’ S; 65° 09LJ 32’’ W, 1160 m asl), located in the province of Jujuy, NW Argentina (Fig. 1). The average annual temperature varies between 18 and 20 °C (Bianchi *et al*. 2005). This area comprises a mixed-use landscape of secondary forests and plantations of exotic *Eucalyptus* sp. (Entrocassi 2015).

**Figure 1.**
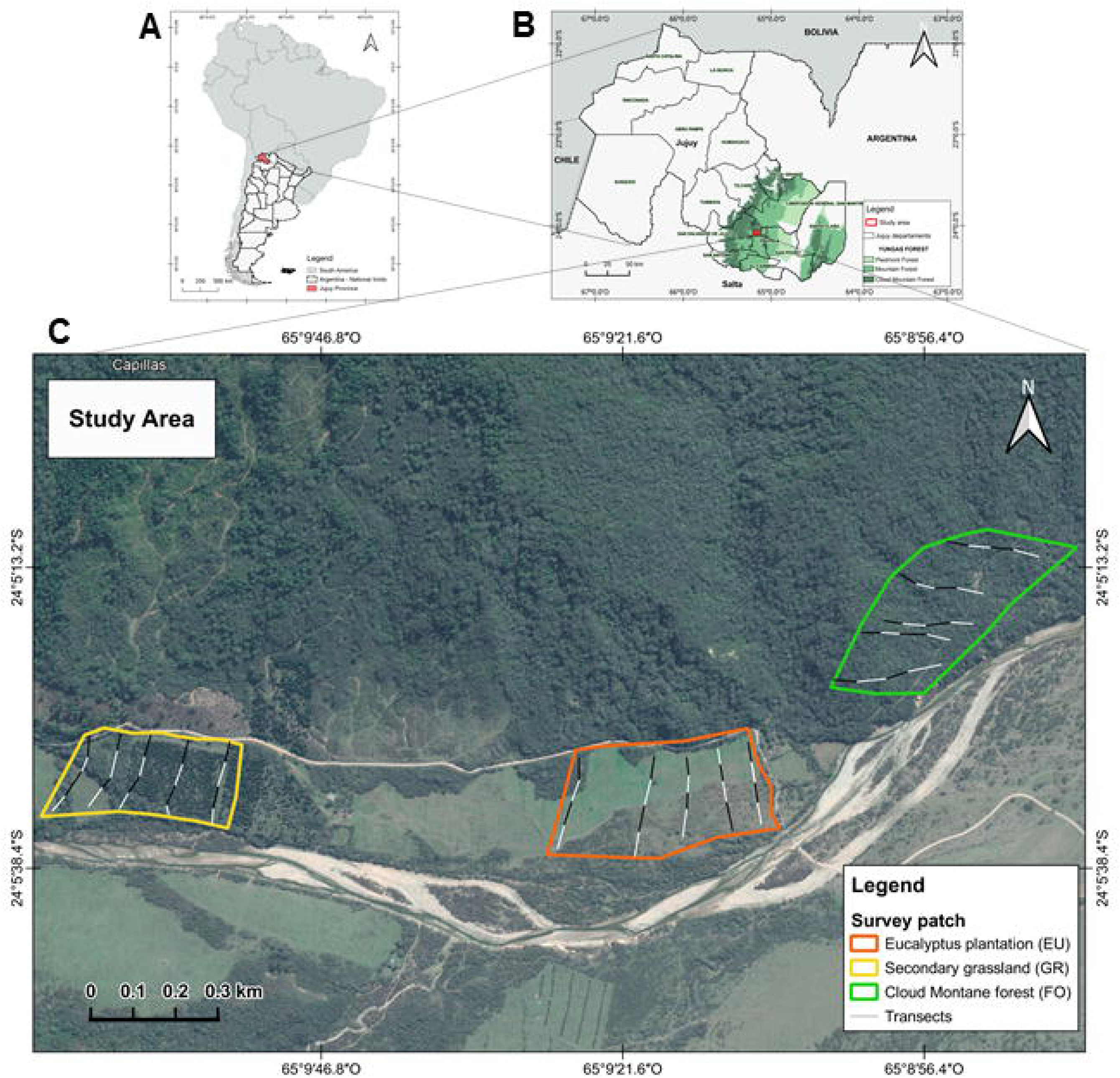
Study area and surveyed patches location at Finca “Las Capillas”, (24° 05LJ 34’’ S; 65° 09LJ 32’’ W, 1160 m asl), Jujuy, NW Argentina. **FO=** Secondary native forest; **EU**= *Eucalyptus* plantation; GR= Secondary grassland.

We selected habitat patches defined by three dominant land-use types: (1) Secondary native forest (FO), covering approximately 0.08 km^2^. The area has previously undergone native wood extraction through selective logging, and it has low-intensity livestock grazing, but at the time of conducting the study, no ongoing extraction was taking place, and logging had been abandoned (pers. obs.). This area was considered a relatively preserved natural forest with minimal land-use change. The area is characterised by trees taller than 15 meters, such as *Tipuana tipu*, *Erythrina falcata*, *Parapiptadenia excelsa*, *Anadenanthera colubrina* var. cebil, *Blepharocalyx salicifolius*, and *Juglans australis*; while *Sebastiania brasiliensis*, *S. commersoniana*, *Allophylus edulis*, *Sapium haematospermum*, *Urera baccifera*, among others, dominate the undergrowth. (Cabrera 1971; Entrocassi 2016). (2) *Eucalyptus plantation* (EU), covering approximately 0.065 km², is intended for timber production, with an estimated age of 4 to 5 years and subject to low-intensity livestock grazing. No evidence of pesticide use was recorded, either at the time of the study or in the past. It has a continuous tree canopy, although within it, there are small remnants of native vegetation and open areas used as livestock pastures, and (3) Secondary grassland (GR), located on flat terrain, featuring secondary grassland with remnants of native forests, and the presence of livestock, covering approximately 0.10 km^2^. This habitat type originated following the primary forest’s clearance and burning. It is dominated by Poaceas and has moderate livestock grazing (Fig. 1).

### Data collection

Butterfly surveys were carried out coinciding with the higher adult butterfly activity in the region and favourable weather conditions, with a similar sampling effort and level of accessibility in all habitats. Sampling took place during three consecutive years (2016-2018) in two different three-day sampling events per habitat patch, one during November/January 2016-2017 and one during April/May 2018 (Autumn). To ensure a comprehensive assessment of diversity, we employed a combination of hand nets and bait trap survey techniques, as both methods are considered complementary (Sparrow *et al*. 1994; Checa *et al*. 2019). In each habitat patch, we placed twenty 50 mts-long transects. We set two Van Someren Rydon bait traps (VSR) (DeVries 1987, 1988; Villareal *et al*. 2004) located at 10 and 40 meters from the start of each transect, accounting for a total of 40 traps per habitat type. The VSRs, manufactured according to the specifications of DeVries (1987, 1988), were slightly modified (i.e. replacing the metal plate of the base with a plastic container). The bait traps, consisting of decaying fruit (banana, apple, peach, mango, pear, depending on seasonal availability in the local market), were suspended 1.5-2.0 m above the ground. The traps were operational for three consecutive days (72 hours) and were visited every 24 hours. Additionally, we conducted entomological net sampling (Villareal *et al*. 2004) along the transects only on the second day after activating the VSR traps, during each sampling event and at each habitat patch. Two researchers sampled flying butterflies using a sweep net for 15 minutes along a standard line transect of 2 m wide (Pollard & Yates 1993), between 11:00 am and 4:00 pm, in coincidence with the peak of butterfly activity (Villareal *et al*. 2004).

Captured butterflies were carefully placed in paper envelopes. Subsequently, specimens were kept in a humid chamber for 24 to 48 hours for later assembly (Villareal *et al*. 2004). All collected material was examined and identified to the species level following Canals (2000, 2003), Lamas (2004), Volkmann & Núñez-Bustos (2010), Klimaitis *et al*. (2018ab), and Guanuco & Martin (2024). In addition, we reviewed the database of Butterflies of America (Warren *et al*. 2023). We followed the taxonomy proposed by Mielke (2008) for the Hesperiidae family, and Lamas (2004) for the remaining Papilionoidea. Approximately three hundred specimens were prepared and deposited at the “Entomological Collection of the Instituto de Biología de la Altura Dra. Liliana Estela Neder (CEINBIAL)”, Universidad Nacional de Jujuy, Jujuy, Argentina.

### Statistical analyses

To evaluate the adequacy of sampling effort at each habitat patch, we employed measures of sample coverage following Chao & Jost (2012). Coverage represents the percentage of individuals in an assemblage that belong to a species represented in the sample; it ranges from 0 to 100% (Chao & Jost 2012). As this value is a measure of inventory completeness, when coverage is close to 100%, the sample is near complete and diversity values (^q^D) can be compared directly (Chao & Jost 2012). We constructed diversity profiles based on Hill numbers (^q^D) to assess the changes in cumulative species diversity among the three habitat patches. We considered *q*-values from 0 to 2 at increments of 0.25 and computed the 95% confidence intervals. Significant differences in species diversity between the two assemblages were inferred when the confidence intervals of the diversity profile curves did not overlap.

We compared species composition among the three habitat patches with the beta-diversity partitioning method following Baselga (2010). This method partitions the total beta diversity into its components of nestedness and species turnover. Nestedness refers to the overlap of species occurrence among habitat patches, indicating that the species composition of a given assemblage is a subset of the species composition of a larger assemblage (Ulrich & Gotelli 2007). Conversely, species turnover refers to the replacement of species in one habitat patch by different ones in another habitat, leading to less frequent or even segregated species occurrence, with many species never co-occurring (Ulrich & Gotelli 2007). To further examine the assemblage structure between habitat patches, we applied a multivariate permutational analysis of variance on Bray-Curtis distances (PERMANOVA; Anderson 2001). Lastly, we assessed changes in community structure with rank-abundance curves based on the log-transformed relative abundances of butterfly species in each habitat patch. These curves visually compare species richness, relative abundance, dominance, and the sequence of each species without losing their identity (Feinsinger 2004). We statistically tested for differences in rank abundance distribution among habitat patches using twoLJsample Kolmogorov–Smirnov tests, under the null hypothesis that two samples can be generated from the same distribution.

All statistical analyses were conducted using R 3.6.1 (R Development Core Team, 2019). The database, R script, and packages are available as supplementary material (Supplemental information, Tables S1 and S2).

## Results

The combined diurnal butterfly collection comprised 2993 individuals, representing an overall diversity of 74 species distributed in six families (Table 1), including Hesperiidae (with two subfamilies: Eudaminae and Pyrginae), Nymphalidae (with nine subfamilies: Apaturinae, Biblidinae, Charaxinae, Danainae, Heliconiinae, Libytheinae, Limenitidinae, Satyrinae, and Nymphalinae), Pieridae (with two subfamilies: Coliadinae and Dismorphiinae), and Lycaenidae, Riodinidae, and Papilionidae, with only one subfamily each (Theclinae, Riodininae, and Papilioninae, respectively).

**Table 1.** List of butterfly species recorded at three habitat patches of Finca “Las Capillas”, (24° 05LJ 34’’ S; 65° 09LJ 32’’ W, 1160 m asl), Jujuy, Argentina. **FO=** Secondary native forest; **EU**= *Eucalyptus* plantation; GR= Secondary grassland. Species for which there is information to consider them exclusive to forest or open areas are highlighted in dark grey (FO) and light grey (GR).

Sample coverage estimates exceeded (95%) for each habitat patch. Diversity profiles based on Hill numbers (CI: 95%) showed that the GR exhibited the highest diversity with 67 species, followed by the EU with 41 species, and FO with 18 species (Table 1, Fig. 2).

**Figure 2.**
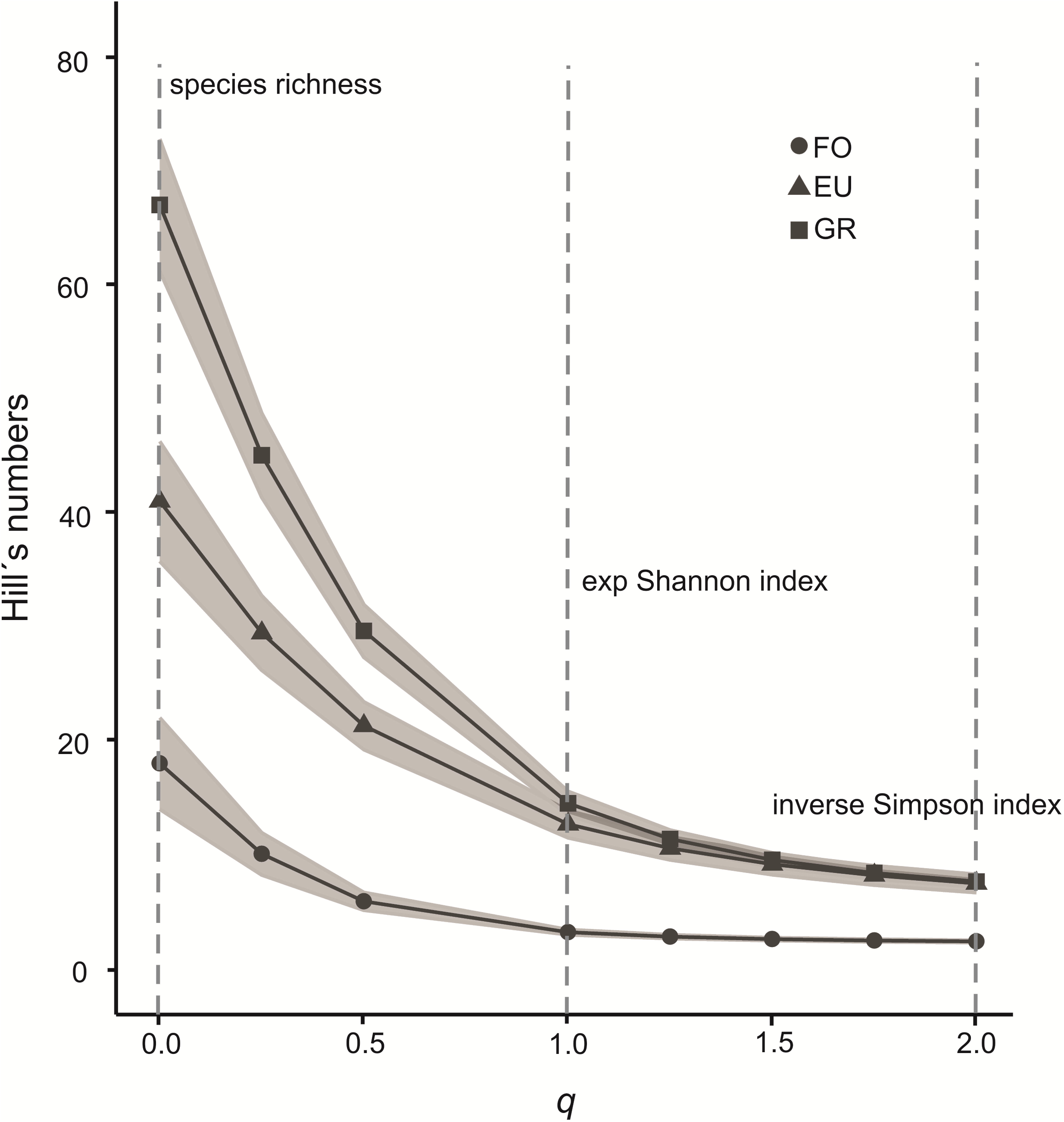
Diversity profiles based on Hill numbers (*^q^*D) for diurnal butterfly assemblages present in three surveyed habitat patches at Finca “Las Capillas”, Jujuy, NW Argentina: **EU**: *Eucalyptus* plantation, **FO**: secondary native forest, and **GR**: Secondary grassland. The x-axis represents the order *q.* Colored ribbons represent the 95% Confidence intervals.

Butterfly species composition differed significantly among the three habitat patches (PERMANOVA, F= 12.43, P <0.001). The FO habitat presented a significant dissimilarity in species composition with both GR and EU habitats, slightly below 60% (PERMANOVA F= 49.52, P =0.003; F= 31.42, P= 0.003, respectively). Observed differences between FO and GR were given mainly by a nestedness process, indicating that the species composition of forest assemblage is a subset of the one found in the grassland (Fig 3). Conversely, beta diversity observed between FO and EU was predominantly driven by species turnover and, to a lesser extent, by a nestedness process, with few species co-occurring in both habitat patches (Fig. 3). Finally, GR and EU presented a relatively lower difference in species composition (Fig. 3), with a statistically significant dissimilarity value of 18% (PERMANOVA F= 8.31, P = 0.003). This dissimilarity was highly influenced by a nestedness process, indicating that the species composition of the *Eucalyptus* sp. assemblage constitutes a subset of the grassland community (Fig. 3).

**Figure 3.**
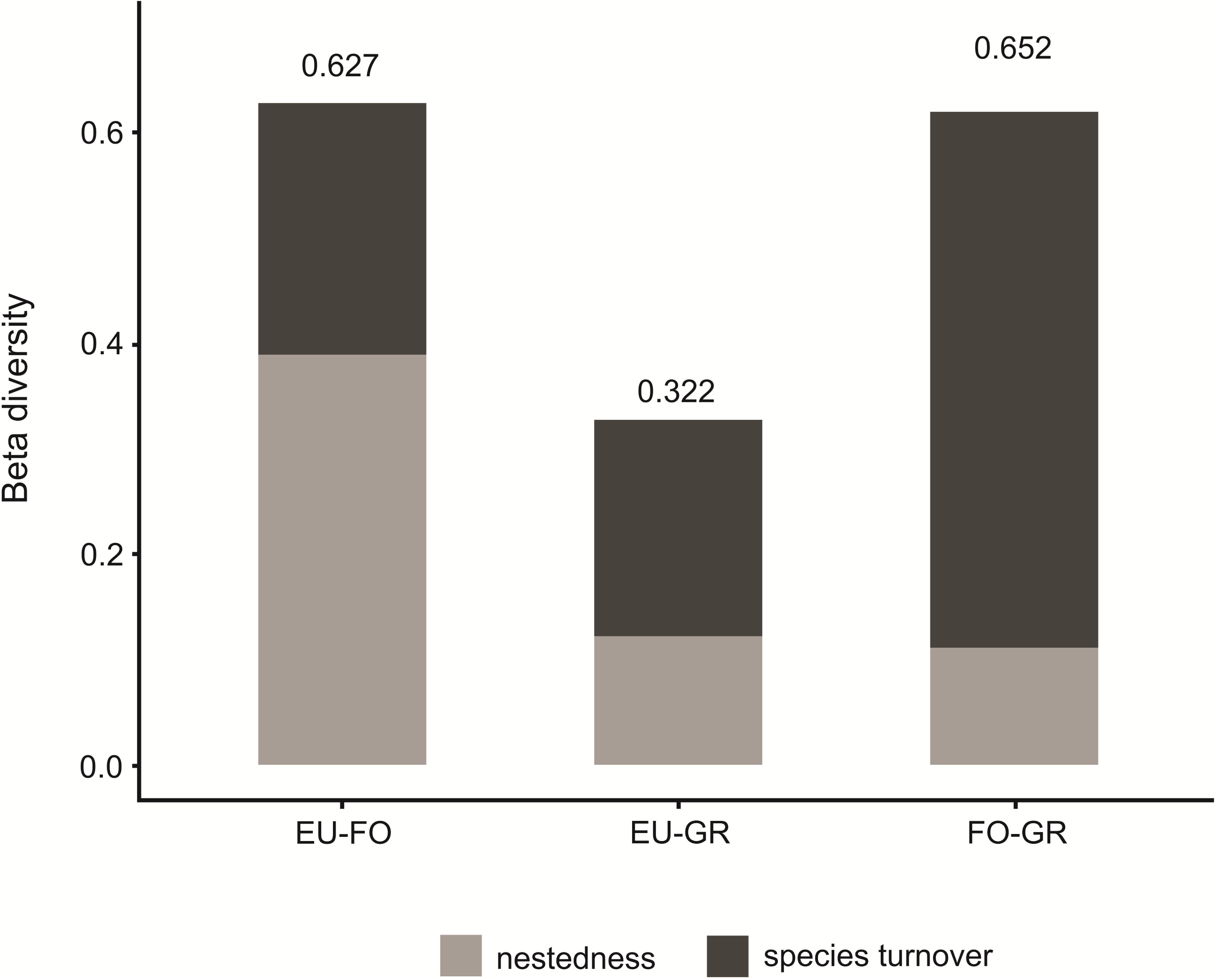
Beta diversity partition into nestedness and species turnover components among the three surveyed habitat patches at Finca “Las Capillas”, Jujuy, NW Argentina: **FO**: secondary native forest, **EU**: *Eucalyptus* plantation, and **GR**: Secondary grassland. Bars correspond to total beta diversity.

The rank-abundance distribution was more even in the grassland assemblage, with three species accounting for over 50% of all individuals registered in this habitat (Fig. 4). In GR, *Biblis hyperia nectanabis* (Fruhstorfer, 1909) (Nymphalidae: Biblidinae) was the most abundant species (25.1%), followed by *Hermeuptychia* sp. Forster, 1964 (Nymphalidae: Satyrinae) (18.3%), and *Cybdelis petronita* Burmeister, 1861 (Nymphalidae: Biblidinae) (14.9%). The EU also presented three species summing more than 50% of the individuals registered in this habitat (Fig. 4): *H.* sp. was again the most abundant species in the assemblage (28.8%), followed by *C. petronita* (12.8%), and *B. hyperia nectanabis* (11.5%).

**Figure 4.**
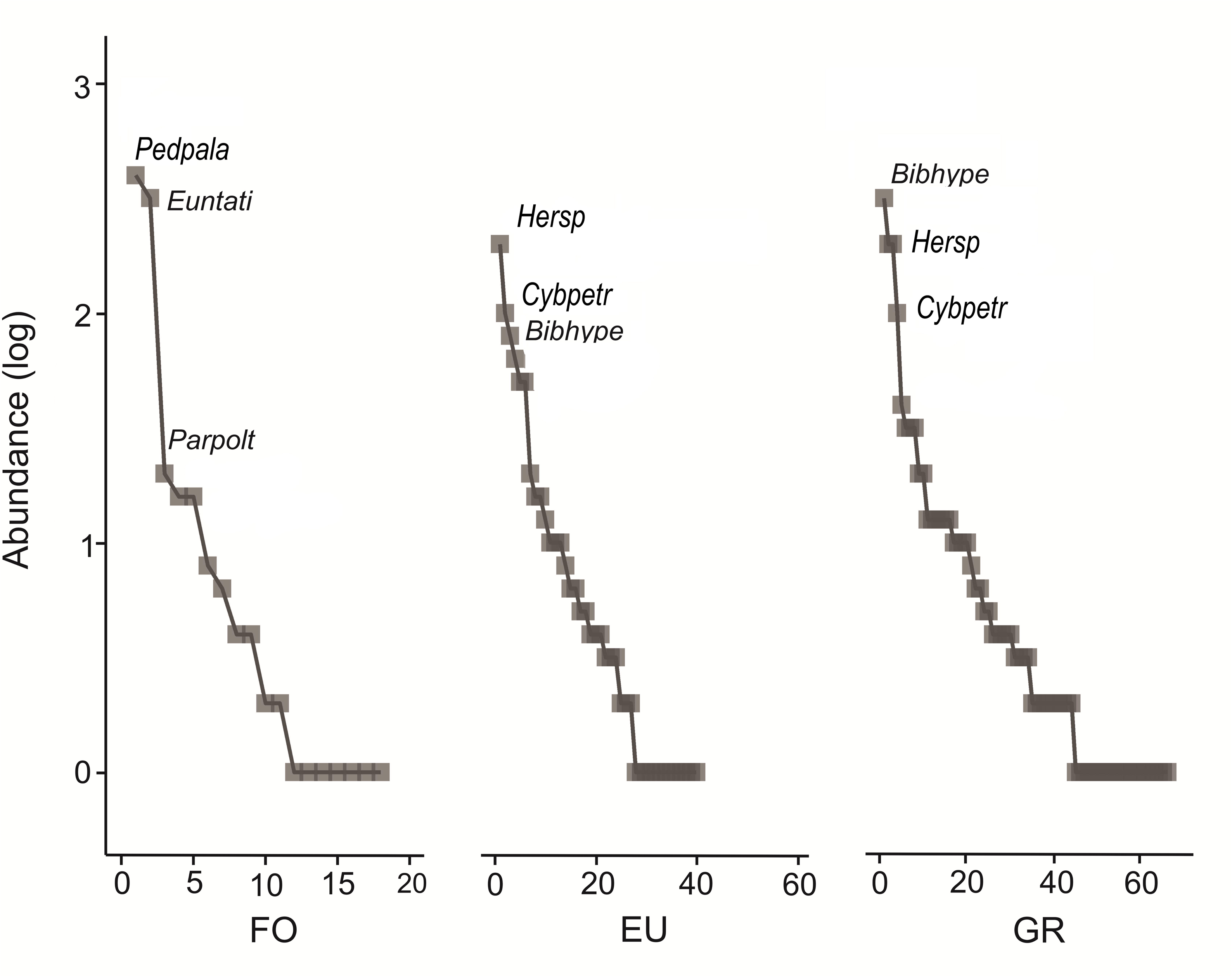
Species rank abundance curves based on the log-transformed relative abundances of butterfly species at each surveyed habitat patch at Finca “Las Capillas”, Jujuy, NW Argentina. **FO**: secondary native forest; **EU**: *Eucalyptus* plantation, and **GR**: Secondary grassland. Only dominant species were identified by a name code (see Abbreviations in Table 1).

In contrast, FO presented two dominant species, which accounted for 90% of the individuals registered for this habitat: *Pedaliodes palaepolis,* (Hewitson, 1878) (Nymphalidae: Satyrinae) (53.4%), and *Eunica tatila bellaria* Fruhstorfer, 1908 (Nymphalidae: Biblidinae) (36.5%) (Fig. 4). However, The Kolmogorov-Smirnov test indicated that the species rank-abundance distribution were similar among the three habitat patches (FO vs. GR: D= 0.15, P =0.70, FO vs. EU: D= 0.18, P= 0.67, and GR vs. EU D= 0.22, P =0.09).

## Discussion

In this study, we documented the diversity of a diurnal butterfly community across habitat patches with different land-use within a sector of the subtropical montane forest of NW Argentina (Yungas ecoregion). Our results suggest that the impacts of land-use change have a strong effect on the diurnal butterfly assemblage of the area. Contrary to our initial expectations, the butterflies assemblage did not show a reduction in species richness with increased land-use change. Instead, we observed the opposite trend, revealing clear differences in alpha diversity among the three habitat patches, where GR emerged as the most diverse habitat, followed by EU and, lastly, FO. The highest diversity registered in GR can be attributed to the structure of the vegetation found in this patch, which includes a small remnant of native forest and open areas with a variety of herbaceous plants. This habitat heterogeneity provides a large amount of resources and niches for several species.

Previous studies established that species belonging to the Satyrinae and Biblidinae subfamilies can display diverse responses to habitat disturbances, with some species being susceptible to such changes, while others became more abundant following alterations (Shahabuddin & Terborgh 1999; Fermon et al. 2005; Barlow et al. 2007ab; Barlow et al. 2009; Uehara-Prado et al. 2007; Ribeiro et al. 2012; Vasconcelos et al. 2019). This aligns with our results, where *Hermeuptychia* sp. (Satyrinae) emerged as the principal satirine species successfully occupying patches of altered habitat compared to the more conserved secondary native forest. Modified environments appeared to be suitable for opportunistic species tolerant to structural changes in the vegetation. The genus *Hermeuptychia* has been geographically expanding in recent years due to its great adaptability, the capacity of larvae to feed on a variety of pioneer plant species belonging to grasses, such as *Poa*, *Paspalum*, and *Stenotaphrum* (Nuñez-Bustos et al. 2010). Therefore, these modified habitats may not only host more satyrin species, but also support a higher abundance per species (Brown 1992). This is consistent with previous studies that identified the genera, *Hermeuptychia* and *Biblis* as pioneering species that first colonize a newly established open area (Ramos 2000). In addition to *B. hyperia nectanabis*, *C. petronita* was also abundant in GR and EU habitat patches, probably due to the association with the type of host plant used during larval development. These butterfly species feed mainly on herbaceous plants of the genus *Tragia* (Freitas et al. 1997; Bauza 2000; Nuñez-Bustos 2010), which are perennials and typically inhabit open areas, such as cultivated fields or stream banks (Romero & Sanguinetti 1989).

Species from the Pieridae family, such as *Abaeis albula albula* (Cramer, 1775), *Teriocolias deva deva* (E. Doubleday, 1847), *Phoebis sennae marcelina* (Linnaeus, 1758), *Pyrisitia nise flóscula* (Cramer 1775), *Anteos clorinde* (Godart 1824), *Abaeis arbela arbela* (Geyer 1832), and *Pseudopieris nehemia prasina*, Hayward, 1949 (Nuñez-Bustos 2010; Klimaitis et al. 2018a; Klimaitis et al. 2018b, Warren et al. 2023), recorded in the habitat patches with the most remarkable land-use intensification, suggest adaptations that enable them to colonize different environments. Their rapid flight, the use of a variety of nutritional plants, and preference for open or edge areas contribute to their capability to occupy extensive modified areas (Ospina López et al. 2010). Other species tolerant to modified environments, such as “jumping” hesperids *Burnsius orcus* (Stoll, 1780), *Urbanus proteus proteus* (Linnaeus, 1758), and certain nymphalids of the subfamily Nymphalinae, represented by heliophilous species of open areas such as *Junonia genoveva hilaris* C. Felder & R. Felder, 1867, *Chlosyne lacinia saundersi* (E. Doubleday, [1847]), *Tegosa claudina* (Eschscholtz, 1821), *Telenassa berenice* (Röber, 1913), *Ortilia gentina* Higgins, 1981, *Ithra ithra* (W. F. Kirby, 1900), *Vanessa braziliensis* (Moore, 1883) and *Vanessa carye* (Hübner, 1812), may colonize different habitats due to their lack of specific habitat requirements (Pastrana et al. 2004; Nuñez-Bustos 2010; Volkmann & Nuñez-Bustos 2010; Klimaitis et al. 2018a; Klimaitis et al. 2018b, Warren et al. 2023). We also recorded species from the Charaxinae subfamily in the modified habitats patches (GR and EU). These lepidopterans, with their large bodies and high mobility, can move across long distances (Barlow et al. 2009; Freire et al. 2021), allowing them to occupy various habitat types.

The absence or reduced abundance of *P. palaepolis* and *E. tatila bellaria* in the GR and EU habitats suggests that these diurnal frugivorous lepidopterans can respond sensitively to habitat disturbances. *P. palaepolis* was recognised as a typical species of montane forest of the Yungas region (F. N. Moschione, pers. comm.). Consequently, the heightened species richness in GR could be attributed to an increase in species tolerant to modified habitats and open areas, resulting in a mechanism of compensatory dynamics, which likely masks the negative effect of land-use change on diversity (Supp & Ernest 2014; Filgueiras *et al*. 2019). Similarly, we considered the EU a heterogeneous habitat despite exhibiting a simplified vegetation structure, including small portions of native forest and open areas, the latter used as cattle pasture. Consequently, species richness in this habitat could also be compensated by common and abundant species, such as *H. sp*., *Yphthimoides celmis*, Godart, (1824), and *Cissia phronius* Godart (1824). These findings were consistent with studies carried out in different regions of Brazil, such as the Amazon Region and the Brazilian Atlantic Forest in Southeast Brazil (Ramos 2000; Ribeiro *et al*. 2012; Vasconcelos *et al*. 2019), where the same genera were registered as abundant and common.

Despite the structural simplification observed in EU, the presence of tree canopy coverage may enable certain species associated with canopy habitats to tolerate land-use changes, explaining the occurrence of species linked to the forest canopy (DeVries 1987). Studies conducted in the Mediterranean regions of Europe show that, despite the challenging arid conditions, butterflies opportunistically use diverse microhabitats such as shaded areas, zones with dense vegetation, and plantations with tree cover. These microhabitats are crucial for protection against predators and also serve as corridors, refuges, and sources of shade (Bonelli *et al*. 2021; Bruschini *et al*. 2024). Although EU did not contribute significantly to the overall biodiversity conservation, it still provides canopy cover, which may offer a relatively safer environment compared to fully deforested areas, where resources are scarce and conditions are more extreme for butterflies. However, this structural simplification could lead to biotic homogenization, often acting as a filter for those species with specific habitat requirements (Bonelli *et al*. 2021), particularly understory butterflies associated with the vertical stratification of a forest, and favouring species capable of using a new spectrum of plants (Ekroos *et al*. 2010; Borschig *et al*. 2013). Furthermore, the absence or decline of certain dominant forest species in EU suggests that some land uses may not adequately preserve local diversity.

Studying species composition through beta diversity measures revealed significant changes in butterfly assemblages, with a concerning reduction of forest-related species. GR and EU, representing higher land-use intensification, exhibited greater similarities to each other, and both differed significantly in community composition from the FO. A nestedness process mainly determined the observed difference in species composition of butterfly assemblages between GR and FO. This suggests that while GR maintains a set of butterfly species relatively similar to the FO, new structural conditions might facilitate the occurrence of a new set of species capable of exploiting highly modified habitats. This may result in a butterfly community increasingly dominated by species adapted to modified environments and open areas, leading to biota homogenization with significant consequences for ecosystem functioning and services of the Yungas forest. On the other hand, the beta diversity observed between FO and EU was mainly driven by a species turnover process.

Comparison of the community structure among the three land-use areas showed not only a significant difference in species composition but also a variation in the dominant species within each habitat type. Species that were abundant in GR and EU, such as *H.* sp. (Satyrinae), *B. hyperia nectanabis* (Biblidinae), and *C. petronita* (Biblidinae), were practically absent in the secondary native forest. Conversely, dominant species in FO, such as *P. palaepolis* (Satyrinae) and *E. tatila bellaria* (Biblidinae), were less abundant or even absent in more modified habitats. *E. tatila bellaria* is characteristic of humid forests associated with habitats of a complex vegetation structure. Its larvae feed on a native tree from the Euphorbiaceae family, specifically the genus *Sebastiania* (Nuñez-Bustos 2010), dominant in the Montane forest strata of the Yungas (Cabrera 1994). Cavanzón-Medrano et al. (2003) showed that *E. tatila* is a neotropical species identified as a bioindicator with significant relevance in undisturbed areas of the Yucatan Peninsula in Mexico. The species predominantly associates with the arboreal stratum, providing essential food resources for the adults (Salazar 2009). Consequently, the dominance of *P. palaepolis* and *E. tatila bellaria* in FO could be considered indicative of low levels of habitat alteration in forest habitats. Montane forest likely serves as a source of host plants, fleshy fruits, and organic matter for larvae and adult stages of these frugivorous species. Our results suggest that, alongside these two species, other forest-dependent butterflies present in the area, like *Taguaiba ypthima* (Hübner, 1821), may also be affected by land-use changes. Its classification as a forest species (DeVries 1987) coincides with our results, given that it was only recorded in FO.

To our knowledge, our work represents the first study on the response of diurnal butterflies to land-use changes in the Southern Andean Yungas ecoregion of Argentina. This research constitutes a first step for future regional-scale studies in this field. Additional studies are needed to assess the impacts of habitat composition and configuration on species diversity in the studied region. Specific habitat requirements and functional traits of the species, such as body size or mobility, may determine whether a particular species benefits from or is adversely affected by intense land-use changes. Also, the dispersal abilities of most species and the distances they travel, crucial for species to cope with alterations in habitat conditions, are still poorly known in the region.

In recent years, concerns have been increasing about declines in overall insect diversity, widely recognized for their roles in ecosystem service provisioning (Hallmann et al., 2021). The greatest threats to conservation of diversity in the Southern Andean Yungas ecoregion are the loss and fragmentation of natural habitats that have occurred relatively recently or are currently taking place (Politi et al. 2021); however, the study of responses of butterfly diversity to land-use change in this region is still incipient. Further research into the mechanisms driving the effects of land-use changes on forest-dependent butterfly diversity could provide insights into which mitigation strategies are most likely to succeed in conserving butterfly diversity in the Yungas of Argentina. Forest-specific studies may identify important threats to the species’ persistence in these natural habitats, and to understand responses of multiple species’ requirements to forest modification. Successful forest management will require an understanding of relationships between habitat characteristics and species diversity, and could help to clarify which kinds of mitigation efforts are most likely to be successful in the Yungas forest and to establish the amount of forest vital to protect even a fraction of the original biodiversity that holds value for local butterfly conservation.

## Acknowledgements

We thank the owners and Juan José Correa, in charge of the “Las Capillas” farm, Sergio José Flores for giving us a place to stay, my friends, family and parents for accompanying me in all the campaigns carried out. Fundación Miguel Lillo partially supported the research of MO (Projects Z-0113-1 and Z-048-1). This project was partially supported by a PIO Grant Conicet #14020140100094CO and a PUE Grant Conicet (#22920170100027CO).

